# BIP4COVID19: Releasing impact measures for articles relevant to COVID-19

**DOI:** 10.1101/2020.04.11.037093

**Authors:** Thanasis Vergoulis, Ilias Kanellos, Serafeim Chatzopoulos, Danae Pla Karidi, Theodore Dalamagas

**Affiliations:** IMSI, “Athena” RC, Athens, Greece; IMSI, “Athena” RC & Univ. of the Peloponnese, Athens, Greece; IMSI, “Athena” RC & NTU Athens, Athens, Greece

## Abstract

Since the beginning of the 2019-20 coronavirus pandemic, a large number of relevant articles has been published or become available in preprint servers. These articles, along with earlier related literature, compose a valuable knowledge base affecting contemporary research studies, or even government actions to limit the spread of the disease and treatment decisions taken by physicians. However, the number of such articles is increasing at an intense rate making the exploration of the relevant literature and the identification of useful knowledge in it challenging. In this work, we describe BIP4COVID19, an open dataset compiled to facilitate the coronavirus-related literature exploration, by providing various indicators of scientific impact for the relevant articles. Additionally, we provide a publicly accessible Web interface on top of our data, allowing the exploration of the publications based on the computed indicators.

## 1 INTRODUCTION

COVID-19 is an infectious disease caused by the coronavirus SARS-CoV-2, which may result, for some cases, in progressing viral pneumonia and multi-organ failure. After its first outbreak in Hubei, a province in China, it subsequently spread to other Chinese provinces and many other countries. On March 11th 2020, the World Health Organisation (WHO) declared the 2019–20 coronavirus outbreak a pandemic. Until the end of February 2021 more than 113 000 000 cases had been recorded in more than 200 countries, counting more than 2 500 000 fatalities. ^1^

At the time of writing, an extensive amount of coronavirus related articles have been published since the virus’ outbreak (indicatively, our collected data contain about 163 986 articles published from 2020 onwards). Taking additionally into account previous literature on coronaviruses and related diseases, it is evident that there is a vast literature on the subject. However, it is critical for researchers or other interested parties (e.g., government officers, physicians) to be able to identify high-impact articles. A variety of impact measures for scientific articles have been proposed in the fields of bibliometrics and scientometrics. Some of them rely on the analysis of the underlying citation network (Kanellos et al., 2019). Other approaches utilise measures commonly known as “altmetrics” (Piwowar, 2013), which analyse data from social media and/or usage analytics in online platforms (e.g., in publishers’ websites). Both approaches have their benefits and shortcomings, each capturing different aspects of an article’s impact. Thus, by considering a wide range of different measures we can better uncover a comprehensive view of each article’s impact.

In this context, the main objective of this work is to produce *BIP4COVID19*, an openly available dataset, which contains a variety of different impact measures calculated for COVID-19-related literature. Four citation-based impact measures (Citation Count, PageRank (Page et al., 1999), RAM (Ghosh et al., 2011), and AttRank (Kanellos et al., 2020)) were chosen to be calculated, as well as an altmetric indicator (Tweet Count). The selected measures were chosen so as to cover different impact aspects of the articles. Furthermore, to select a representative set of publications, we rely on two open datasets of COVID-19-related articles: the CORD-19 (L. L. Wang et al., 2020) and the LitCovid (Q. Chen et al., 2020a,b) datasets. BIP4COVID19 data are updated on a regular basis and are openly available on Zenodo ^2^ (Vergoulis, Kanellos, Chatzopoulos, et al., 2021). Since its initial launch in March 2020, 43 versions of BIP4COVID19 have been released, with Zenodo recording more than 90, 000 views and 10, 000 downloads. Apart from the dataset itself, we have developed and released a freely accessible Web-interface3, which functions as a search engine on COVID-19 related literature. This search engine exploits the calculated measures to offer relevant functionalities (e.g., impact-based ranking for results) that facilitate the exploration of the COVID-19 related literature.

## 2 MATERIALS AND METHODS

BIP4COVID19 is a regularly updated dataset. Data production and update is based on the semi-automatic workflow presented in Figure 1. In the following subsections the major processes involved are elaborated.

**Fig. 1.**
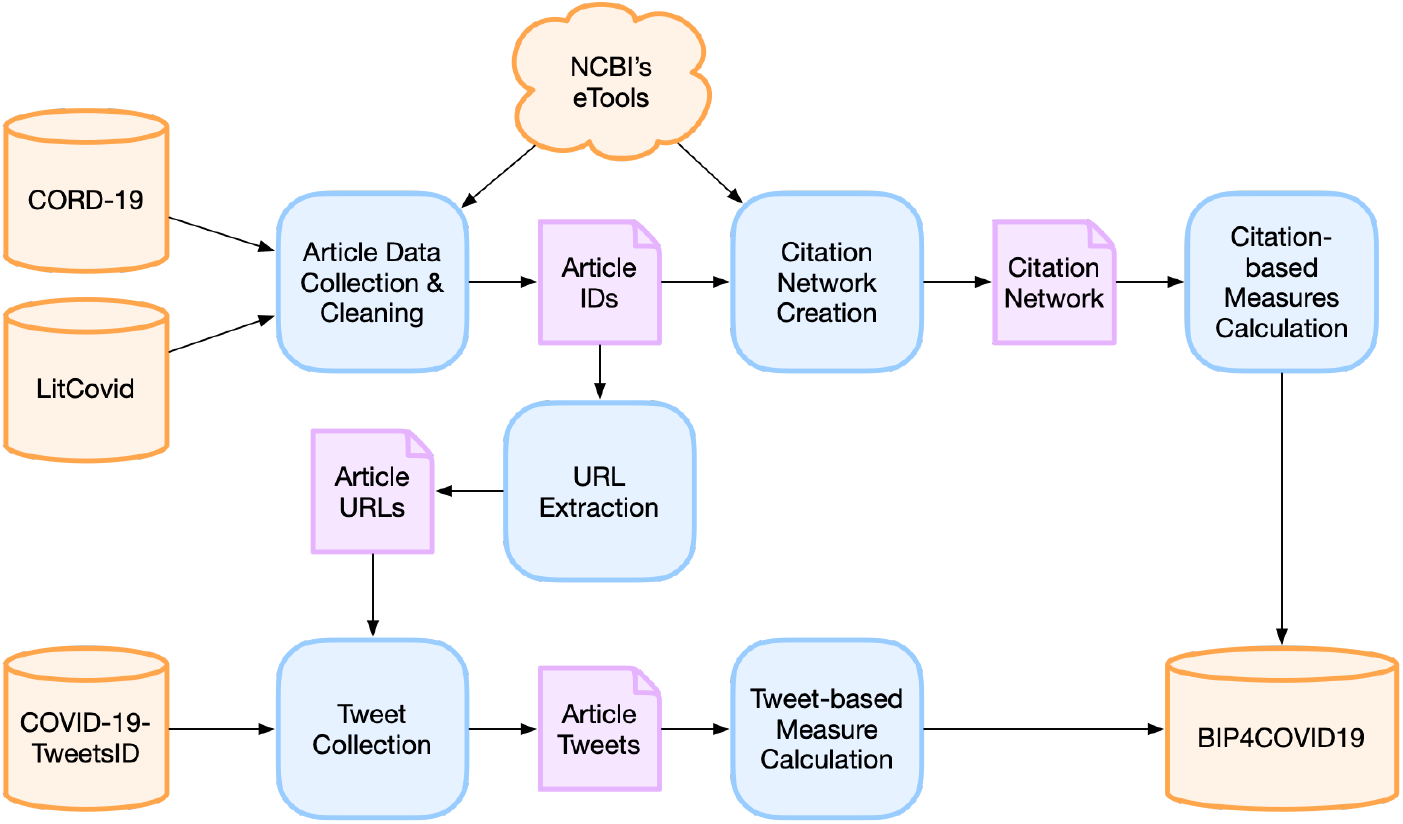
The data update workflow of BIP4COVID19

### 2.1 Article Data Collection and Cleaning

The list of COVID-19-related articles is created based on two main data sources: the *CORD-19* ^4^ Open Research Dataset (L. L. Wang et al., 2020), provided by the Allen Institute for AI, and the *LitCovid* ^5^ collection (Q. Chen et al., 2020a) provided by the NLM/NCBI BioNLP Research Group. CORD-19 offers a full-text corpus of more than 467 000 articles on coronavirus and COVID-19, collected based on articles that contain a set of COVID-19 related keywords from PMC, arXiv, biorXiv, and medRxiv and the further addition of a set of publications on the novel coronavirus, maintained by the WHO. LitCovid, is a curated dataset which currently contains more than 103 000 papers on the novel coronavirus.

The contents of the previous datasets are integrated and cleaned. During this process, the eSummary tool ^6^ from NCBI’s eTool suite is utilised to collect extra metadata for each publication using the corresponding PubMed or PubMed Central identifiers (pmid and pmcid, respectively), where available. The collected metadata are semi-automatically processed to remove duplicate records. The resulting dataset contains one entry for each distinct article. Each entry contains the pmid, the DOI, the pmcid, and the publication year of the corresponding article. This information is the minimum required for the calculation of the selected impact measures.

### 2.2 Calculation of Citation-based Measures

A prerequisite for calculating the citation-based impact measures of the collected articles, is the compilation of their citation network, i.e., the network which has articles as nodes and citations between them as directed edges. The citations of the articles required to construct this network are gathered using NCBI’s eLink tool. The tool returns for a given article the identifiers (pmids/pmcids) of all articles that cite, or are cited by it. Four citation-based impact measures are calculated for each article using the constructed network:

- the *Citation Count* score, which sums all citations received by the article.
- the PageRank (Page et al., 1999) scores, based on the well-known Web page ranking algorithm, which has been useful in ranking papers in citation networks (e.g., (P. Chen et al., 2007)).
- the RAM (Ghosh et al., 2011) scores, which are weighted citation counts, where recent citations are considered as more important.
- the AttRank (Kanellos et al., 2020) scores, which are based on a PageRank variant that focuses on overcoming PageRank’s bias against recent papers.

Citation Count was selected because it is the most widely known measure; the other three were selected based on the results of a recent experimental study (Kanellos et al., 2019) and subsequent work (Kanellos et al., 2020), which found them to perform best in capturing the overall and the current impact of an article (i.e., its “influence” and its “popularity”), respectively. In particular, PageRank evaluates the overall impact of articles by differentiating their citations based on the importance of the articles making them. However, it is biased against recent articles that haven’t accumulated many citations yet, but may be the current focus of the research community. RAM alleviates this issue by considering recent citations as being more important, while AttRank modifies PageRank so that it promoted recently published or recently cited papers, an approach which has been found to perform more effectively in ranking papers based on their current impact (Kanellos et al., 2020). Based on the previous discussion, we suggest that the PageRank score should be preferred to capture influence, while the AttRank score should be preferable for applications that focus on popularity.

### 2.3 Calculation of Tweet-based measure

In addition to the citation-based measures, for each article, the number of recent tweet posts mentioning it is calculated, as well. This is considered a measure of its social media attention. The *COVID-19-TweetIDs* ^7^ dataset (E. Chen et al., n.d.) is used for the collection of COVID-19-relevant tweets. This dataset contains a collection of tweet IDs, each of them published by one of 9 predetermined Twitter accounts (e.g., @WHO) and containing at least one out of 71 predefined coronavirus-related keywords (e.g., “Coronavirus”, “covid19”, etc). At the time of writing, a subset of this dataset containing tweets posted from January 30 ^*th*^2021 to February 5 ^*th*^2021 (25 852 753 unique tweet IDs) have been integrated in BIP4COVID19. The corresponding Tweet objects were collected using the Twitter API. The result was a collection of 20 529 951 tweet objects. The difference between the number of IDs and hydrated objects is due to facts, such as the deletion of tweets in the meantime, which makes some tweets impossible to retrieve.

To find those tweets which are related to the articles in our database, we rely on the URLs of the articles in doi.org, PubMed, and PMC. These URLs are easily produced based on the corresponding identifiers. In addition, when possible, the corresponding page in the publisher’s website is also retrieved based on the doi.org redirection. After the collection of the URLs of all articles, the number of appearances of the URLs related to each one are produced. However, since the Twitter API returns either shortened or not fully expanded URLs, the fully expanded URLs are collected using the unshrtn ^8^ library.

## 3. RESULTS

### 3.1 Dataset Details

The BIP4COVID19 dataset, produced by the previously described workflow, is openly available on Zenodo (Vergoulis, Kanellos, Chatzopoulos, et al., 2021), under the Creative Commons Attribution 4.0 International license. Its first version was released on March 21 ^*st*^ 2020. At the time of publication, the forty third release of this dataset (v36) is available ^9^, counting 255 914 records in total. Of these, 229 043 correspond to entries in PubMed, 185 909 to entries in PMC, while 248 480 have an associated DOI. All publications included were published from 1825 to 2021. The distribution of publication years of the articles recorded in the dataset is illustrated in Figure 2. 163 986 of these articles were published in 2020, i.e., after the coronavirus pandemic outbreak, while 91 928 were published from 1825 to 2019. Moreover, the number of articles per venue for the top 30 venues (in terms of relevant articles published) are presented in Figure 3.

**Fig. 2.**
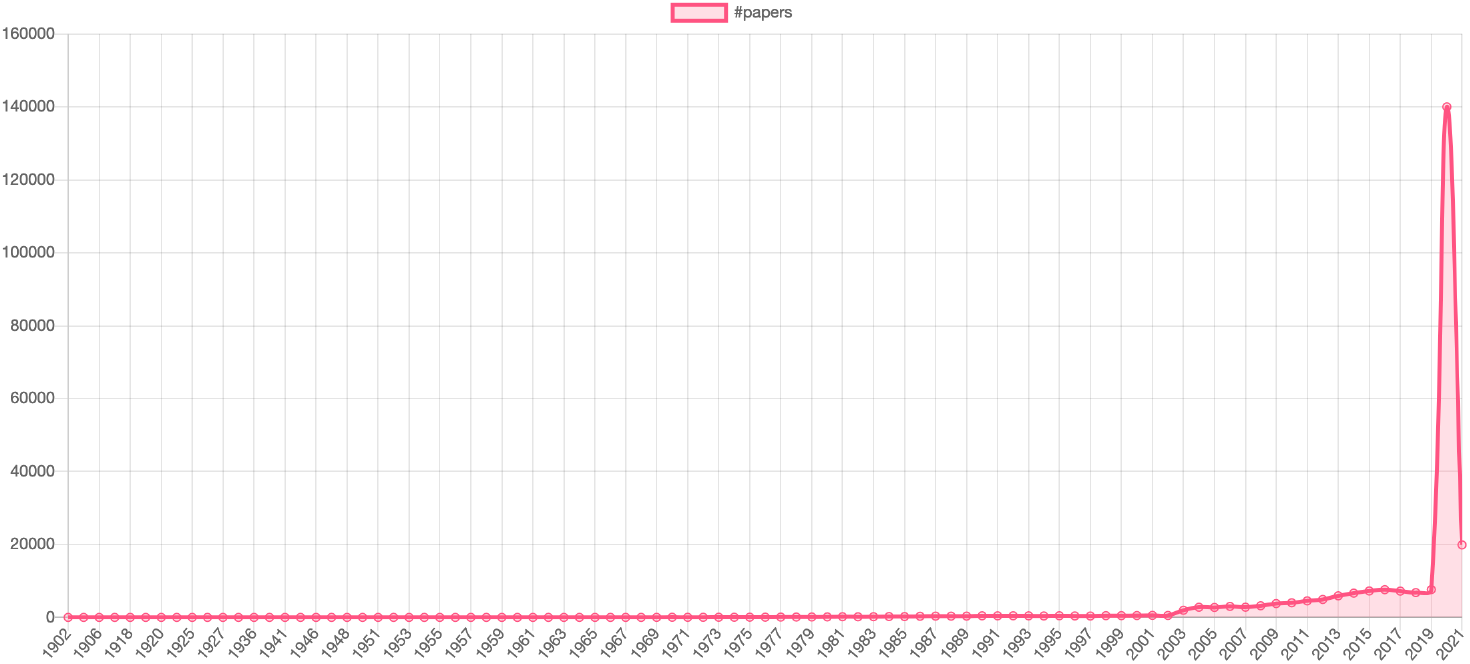
COVID-19-related articles per year. Papers published before 1902 are excluded for presentation reasons.

**Fig. 3.**
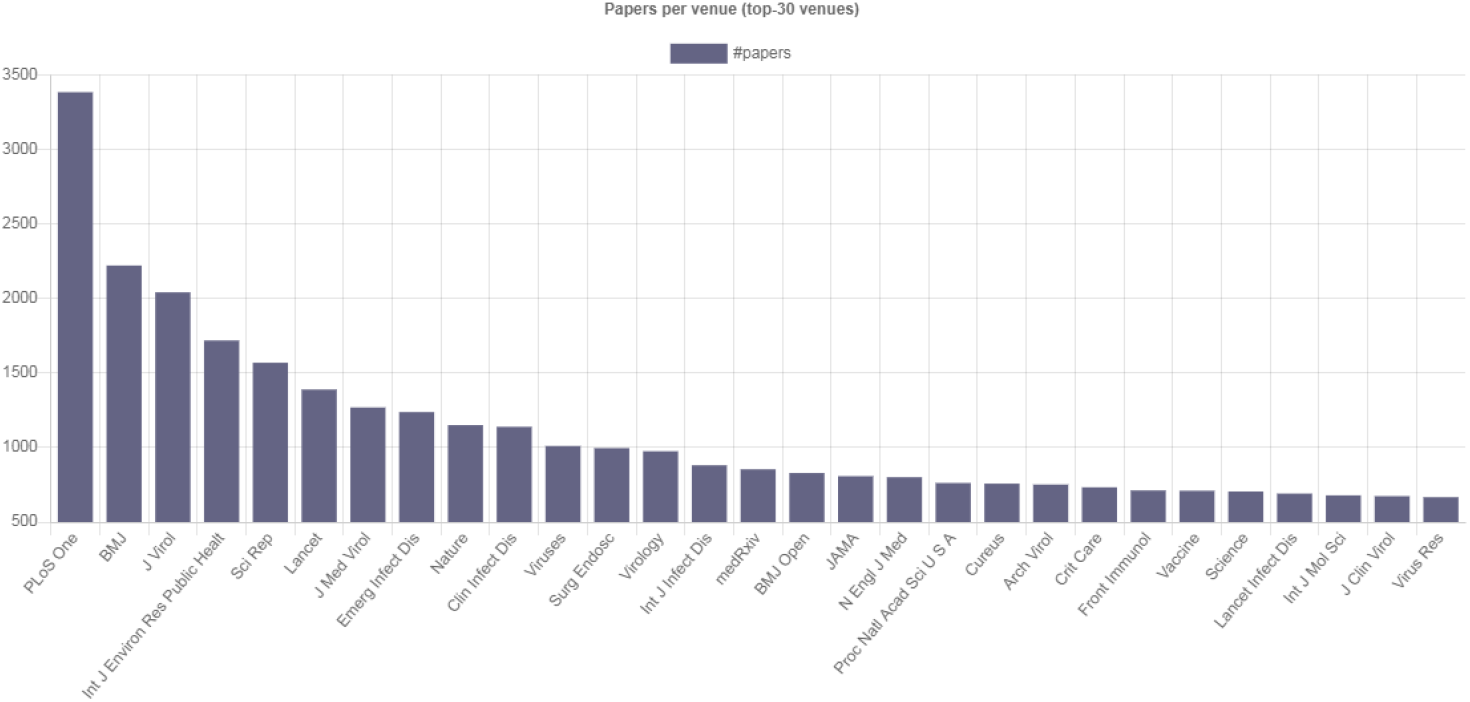
Top 30 venues in terms of published COVID-19-related articles.

The BIP4COVID19 dataset is comprised of five files in tab separated (TSV) format. The files contain identical information, however, in each of them, the records are ordered based on a different impact measure. It is worth mentioning that, at the time of writing, the BIP4COVID19 dataset has already been downloaded more than 10 000 times (according to the metrics provided from Zenodo for all its versions), while it appears as the most viewed dataset (among 423 datasets) in the “Coronavirus Disease Research Community - COVID-19” collection in Zenodo.

### 3.2 Web-based Search Engine

A Web interface has been developed on top of the BIP4COVID19 data. Its aim is to facilitate the exploration of COVID-19-related literature. Apart from a basic keyword search functionality, the option to order articles according to different impact measures is provided. This is expected to be useful since users can better prioritise their reading based on their needs. For example, a user that wants to delve into the background knowledge about a particular COVID-19-related sub-topic could select to order the articles based on their influence. On the other hand, another user that needs to get an overview of the latest trends in the same topic, could select to order the articles based on their popularity. It is worth mentioning that, to avoid user confusion, the Web Interface incorporates only one influence measure (PageRank) and one popularity measure (AttRank); the selection was based on their ranking effectiveness based on previous studies (Kanellos et al., 2019, 2020). Finally, the Web user interface takes advantage of the Mendeley API ^10^ to also display the current number of Mendeley readers, as an extra altmetric measure.

The information shown to users, per publication, includes its title, venue, year, and the source dataset where it was found (see Figure 4). Moreover, each result is accompanied by color coded icons that denote the publication’s importance based on each calculated impact measure. In this way, users can easily get a quick insight about the different impact aspects of each article. The tooltips of these icons provide the exact scores for each measure. Each publication title functions as a link to the corresponding article’s entry in its publisher’s website, or to Pubmed. Additionally, if an article has been retracted, the corresponding entry in the results page is accordingly flagged (we collect this information from NCBI’s esummary tool ^11^, which provides summary information for publications, including their possible retraction status, given their identifier in pubmed). Finally, a page containing various summary statistics about the BIP4COVID19 data is provided. This page contains various charts that visualise, for example, the number of articles per year, or the number of articles that have substantial impact based on each of the provided impact measures, per year.

**Fig. 4.**
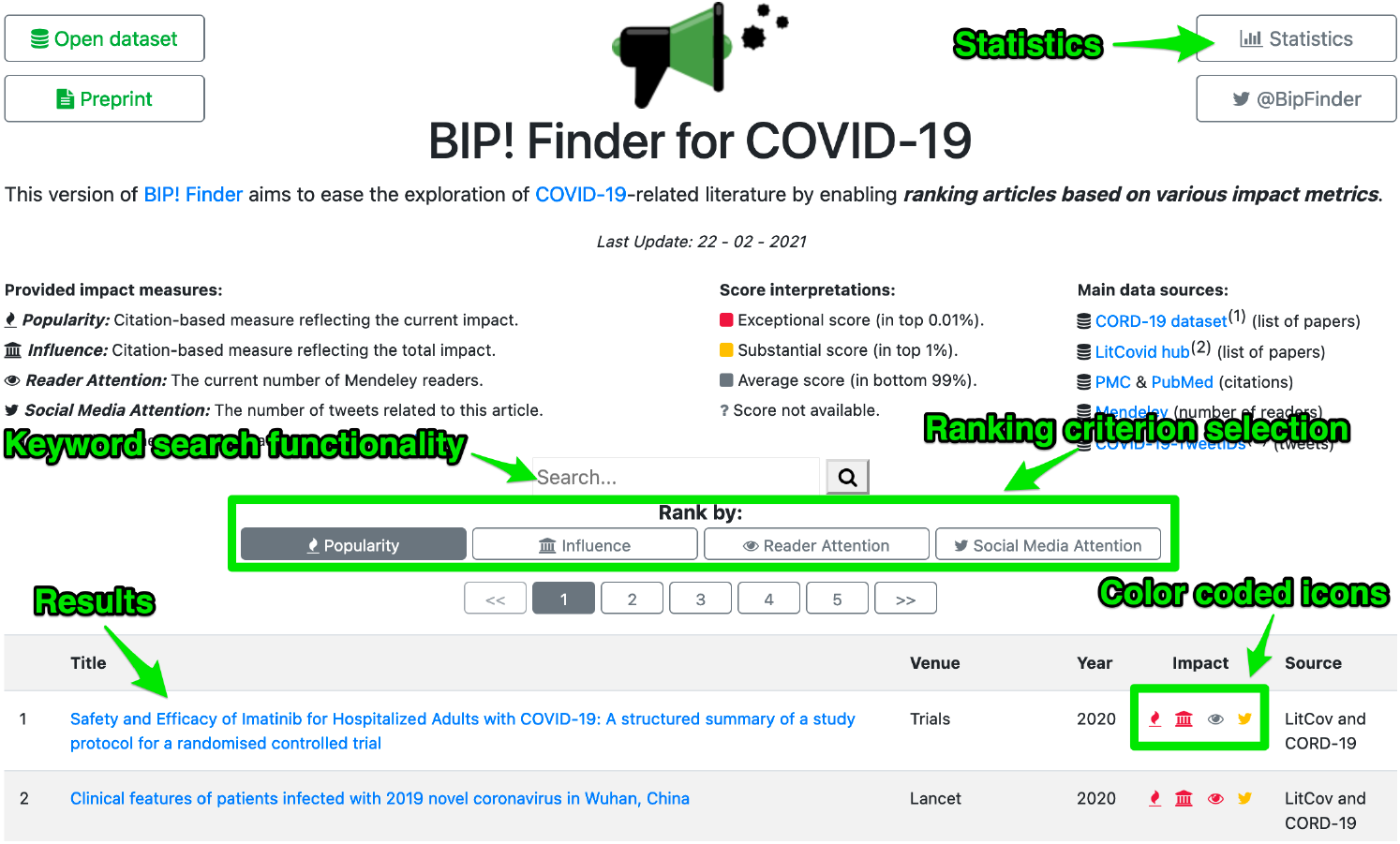
A screenshot of the BIP4COVID19 Web interface.

## 4. DISCUSSION

### 4.1 Dataset Analytics

In Figures 5-7 we present various analytics on the BIP4COVID19 dataset. Figure 5 presents the timeline of the dataset size (in thousands of papers recorded). Since its first versions the dataset has shown a steady linear increase in the number of papers recorded, which follows the corresponding increase in the size of the CORD-19 and litcovid datasets.

**Fig. 5.**
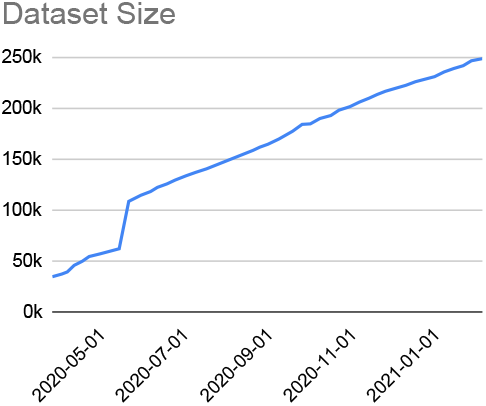
BIP4COVID19 dataset size timeline.

In Figure 6 we examine the overall correlation of the influence- and popularity-based rankings. The rankings examined are based on PageRank for influence and RAM for popularity. ^12^ We measure correlation using Spearman’s *ρ*, which ranges in[−1, 1] where *ρ* = −1 denotes perfect correlation and *ρ* = −1 denotes perfect inverse correlation. From Figure 6, we observe that the popularity- and influence-based rankings are relatively highly correlated (*ρ*∼ 0.7 throughout the dataset’s timeline). While popularity and influence are highly correlated, they do not however correlate perfectly. Figure 7 shows the timeline of the overlapping top-100 ranked papers, based on popularity and influence. We observe that for all versions of the BIP4COVID19 dataset the overlap of top-ranking papers is less than 50%, while for early versions there was no overlap. These results complement Figure 6, in highlighting that popularity- and influence-based rankings, while correlated, do indeed constitute rankings with distinct semantics, which correspond to different types of impact.

**Fig. 6.**
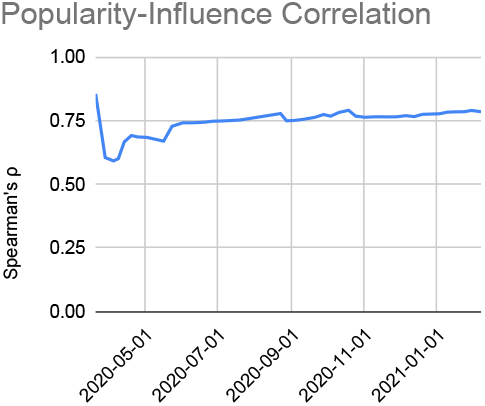
Popularity-Influence rank- ing correlation timeline.

**Fig. 7.**
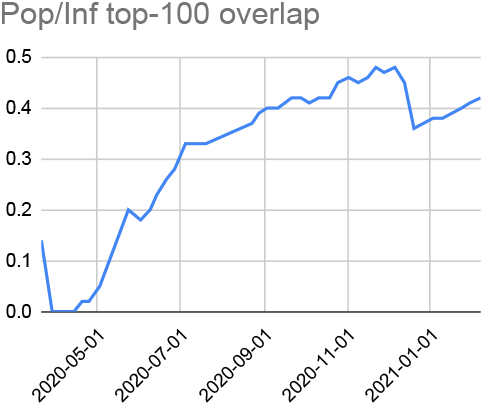
Timeline of top-100 ranking papers by Influence and Popularity.

### 4.2 Ensuring Data Integrity

To ensure the proper integration and cleaning of the CORD-19 and LitCovid datasets, we rely on NCBI’s eTool suite. In particular, we collect pmids and pmcids from both datasets and use them as queries to gather each article’s metadata. After cleaning the article title (e.g., removing special characters) we automatically identify duplicates by comparing each record’s complete content and eliminate them. Finally, manual inspection is performed to produce the correct metadata for a limited number of duplicates that remain (e.g., duplicate records containing the title of the same publication in two different languages).

Further, to guarantee the correctness of the compiled citation graph we apply the following procedures. After gathering all citing - cited records using NCBI’s eTools, those that include identifiers not found in the source data are removed. Since many citing - cited pairs may have been found both with pmids and pmcids, the resulting data may still contain duplicate records. These records are removed, after mapping all pmids/pmcids to custom identifiers, with pmid-pmcid pairs that refer to the same article being mapped to the same identifier. The final resulting citation graph is based on these mapped identifiers. As an extra cleaning step, any links in the graph that denote citations to articles published at a later time than the citing article are removed.^13^

To ensure that we retrieve a set of tweets about each article that is as comprehensive as possible, we collect not only the URLs in doi.org, Pubmed, and PMC, but also the URL to the article in its publisher’s website, where possible. These latter URLs are very important, since they are widely used in tweets. To collect them we utilize doi.org redirections. To avoid incorrect tweet counts due to duplicate tweets, we used a simple deduplication process after the Tweet object retrieval. Moreover, the use of the unshrtn library to expand the short URLs from tweet texts ensures that our measurements derive from all available URL instances of each publication record, no matter how they were shortened by users or Twitter.

### 4.3 Limitations

The following limitations should be taken into consideration with respect to the data: while we take effort to include as many articles as possible, there are many cases where our source data do not provide any pmids or pmcids. As a consequence, no data for these articles are collected and they are not included in the BIP4COVID19 dataset. Furthermore, with respect to the calculated impact scores, it should be noted that the citation analysis we conduct is applied on the citation graph formed by citations *from* and *to* collected publications only, i.e., our analyses are not based on pubmed’s complete citation graph, but on a COVID-19-related subgraph. Consequently, the relative scores of publications may differ from those calculated on the complete PubMed data. Finally, regarding the tweet-based analysis, since our data come from the COVID-19-TweetIDs dataset which only tracks tweets from a predefined set of accounts and which is based on a particular set of COVD-19-related keywords, the measured number of tweets is only based on a subset of the complete COVID-19-related tweets.

### 4.4 Usage Notes

Our data are available in files following TSV format, allowing easy import to various database management systems and can be conveniently opened and edited by any text editor, or spreadsheet software. We have been regularly updating the BIP4COVID19 data since March 2020, and we plan to continue providing regular updates, incorporating any additions and changes from our source datasets. Additionally, we plan to incorporate any further sources on coronavirus related literature that may be released and which will index the literature based on pmids and/or pmcids.

The contents of the BIP4COVID19 dataset may be used to support multiple interesting applications. For instance, the calculated scores for each impact measure could be used to rank articles based on their impact to help researchers prioritise their reading. In fact, we used our data to implement such a demo as previously described. Additionally, the impact scores may be useful for monitoring the research output impact of particular sub-topics or as features in machine learning applications that apply data mining on publications related to coronavirus. Finally, the set of impact measures provided may be useful as a basis for various scientometric analyses, e.g., for studying the relations between the various impact measures, studying the patterns connecting them, or gaining other insights hidden in these data.

## 5. RELATED WORK

### 5.1. Studies of the COVID-19 Literature

During the last months, various works studying COVID-19-related scientific literature have been published. The authors of (Kousha & Thelwall, 2020) evaluate the coverage of various scholarly data sources regarding COVID-19-related articles during the time period between March 21th, 2020 and April 18th, 2020. They also report how many articles received a significant number of citations or a substantial mass and social media attention, during the same period. An interesting finding of their analysis is that, for COVID-19 papers, the convergence between citation counts and social media attention appears to be high, something that has not been observed for other fields of study. The authors of (Horbach, 2020) investigate whether medical journals have managed to accelerate their publication processes for COVID-19-related articles. They studied the duration time of the publication process of 14 medical journals before and after the start of the COVID-19 pandemic and found that the time between submission and publication for COVID-19-related articles was 49% smaller, on average, than the corresponding times of other articles during the same period or in the past.

In (Thelwall, 2020), the authors use Mendeley reader counts to reveal that studies analysing SARS and MERS data to provide useful intuition about COVID-19 have gathered more academic interest than primary studies of SARS and MERS, on average. Finally, the authors of (Colavizza, 2020) focus on COVID-19-related Wikipedia pages and investigate the pace with which the results of new research are incorporated into them and which proportion of the relevant literature is covered into Wikipedia pages.

Although we provide a small set of statistics about the recorded COVID-19-related articles (see Section 4.1), the main focus of this work is to provide an open, frequently updated dataset of impact measures for the relevant literature. Nevertheless, the statistics we provide are complementary to those given by the aforementioned studies, since we focus on different research questions.

### 5.2 Available Datasets

A number of openly available datasets have been released and actively maintained to facilitate the COVID-19 related literature exploration and research. The most popular example is the COVID-19 Open Research Dataset (CORD-19) (L. L. Wang et al., 2020). CORD-19 is a frequently updated dataset of publications, which, apart from COVID-19 research articles, contains current and past research related to the coronavirus family of viruses. It combines articles from different sources (PMC, arXiv, biorXiv, medRxiv and the World Health Organization) and offers deduplicated metadata and a full text corpus for a large number of articles. In another line of work, the authors of (Rohatgi et al., 2020) further enhanced a version of CORD-19 with additional abstracts, tables and figures also providing *COVIDSeer*, a keyword search engine on top of the enhanced dataset (see Section 5.3). Another very popular example is LitCovid (Q. Chen et al., 2020a,b), an openly available curated literature resource of COVID-19 related articles, further categorised by research topics and geographic locations for improved access. Moreover, there are various attempts for the creation of knowledge graphs that capture information contained in the COVID-19 related literature (Wise et al., 2020; Q. Wang et al., 2020). Finally, there are also various large-scale datasets collecting COVID-19 related tweets (E. Chen et al., n.d.; Banda et al., 2020), while Twitter itself released its own API endpoint to provide real-time access to COVID-19 related tweets.

Our own dataset, BIP4COVID19, focuses on providing a range of different impact measures on COVID-19 related publications. This line of work aims at providing a complete picture about a publication’s impact, in line with the best practices in research assessment. In fact, our work is a COVID-19-specific version of BIP! DB (Vergoulis, Kanellos, Atzori, et al., 2021). To the best of our knowledge, BIP4COVID19 is the only dataset of this scale to provide a fair range of different impact measures on COVID-19 related literature.

### 5.3 COVID-19 Search Services

Following the beginning of the COVID-19 outbreak, due to the extremely large interest in COVID- 19-related scientific articles, various scholarly search engines, which are tailor-made for the COVID-19-related literature have been developed. First of all, the teams that developed and maintain the major literature datasets provide their own search engines: Allen Institute for AI released *SciSight* (Hope et al., 2020), a tool for exploring the CORD-19 data, while *LitCovid* (Q. Chen et al., 2020a,b) provides a search engine featuring basic functionalities (e.g., keyword search, facets). *COVIDScholar* (Trewartha et al., 2020) is another search engine that uses natural language processing (NLP) to power search on a set of research papers related to COVID-19. *iSearch COVID-19 Portfolio* ^14^ is a tool that provides search functionality and faceting for COVID-19 articles. *KDCovid* ^15^ is a tool that retrieves papers by measuring similarity of queries to sentences in the full text of papers in CORD-19 corpus using a similarity metric derived from BioSentVec. *COVIDSeer* (Rohatgi et al., 2020) is another tool that was built on top of a CORD-19 extension and which provides keyword filtering and paper recommendation functionalities. *Vilokana* (Panja et al., 2020) is a semantic search engine on top of the CORD-19 dataset that exploits a set of modified TFIDF features and cosine similarity with ontology maps to provide semantic search functionalities. *CAiRE-COVID* (Su et al., 2020) is a tool that answers priority questions and summarizing salient question-related information. It combines information extraction with state-of-the-art QA and query-focused multi-document summarisation techniques, selecting and highlighting evidence snippets from existing literature based on a given query. Finally, *AWS CORD-19 Search (ACS)* (Bhatia et al., 2020) is a COVID-19 specific, ML-based search engine that supports natural language based searches providing document ranking, passage ranking, question answering and topic classification functionalities.

To the best of our knowledge, apart from a couple of engines that display citation count scores foreach article in the result list, or use them to rank the results, most of them do not leverage impact measures. Incorporating the impact measures, which are openly available through the BIP4COVID19 dataset, may unlock various opportunities towards providing advanced search, filtering, monitoring, and other valuable features. Recognising these opportunities, we have developed our own prototype search engine (see Section 3.2). It is worth mentioning that this tool is a COVID-19-specific variant of *BIP! Finder* (Vergoulis et al., 2019).

## 6 CONCLUSION

We presented BIP4COVID19, an openly available dataset, providing impact scores for coronavirus related scientific publications. Our dataset can be potentially useful for many applications like assisting researchers to prioritize their reading of COVID-19 related papers, providing study material for scientometricians, etc. We have additionally built on our dataset, providing a Web-based article search engine which exploits the calculated impact measures to offer relevant functionalities, such as impact-based ranking of the keyword-search results.

## ACKNOWLEDGMENTS

We acknowledge support of this work by the project “Moving from Big Data Management to Data Science” (MIS 5002437/3) which is implemented under the Action “Reinforcement of the Research and Innovation Infrastructure”, funded by the Operational Programme “Competitiveness, Entrepreneurship and Innovation” (NSRF 2014-2020) and co-financed by Greece and the European Union (European Regional Development Fund).

## Footnotes

1 https://covid19.who.int/

2 https://zenodo.org/record/4555117

3 https://bip.covid19.athenarc.gr/

4 https://pages.semanticscholar.org/coronavirus-research

5 https://www.ncbi.nlm.nih.gov/research/coronavirus/

6 https://www.ncbi.nlm.nih.gov/books/NBK25500/

7 https://github.com/echen102/COVID-19-TweetIDs

8 https://github.com/docnow/unshrtn

9 The dataset is updated on a weekly basis.

10 Mendeley APIhttps://dev.mendeley.com/

11 https://www.ncbi.nlm.nih.gov/books/NBK25499/

12 We chose these measures because citation counts and AttRank scores have only been included in the most recent versions.

13 Such references to future articles are often observed in citation data due to various reasons. Hence, a common practice is to remove them (Ghosh et al., 2011).

14 https://icite.od.nih.gov/covid19/search/

15 http://kdcovid.nl/

## References

Banda, J. M., Tekumalla, R., Wang, G., Yu, J., Liu, T., Ding, Y., … Chowell, G. (2020). A large-scale covid-19 twitter chatter dataset for open scientific research – an international collaboration.

Bhatia, P., Arumae, K., Pourdamghani, N., Deshpande, S., Snively, B., Mona, M., … Kass-Hout, T. A. (2020). AWS cord19-search: A scientific literature search engine for COVID-19. CoRR, abs/2007.09186. Retrieved from https://arxiv.org/abs/2007.09186

Chen, E., Lerman, K., & Ferrara, E. (n.d.). Covid-19: The first public coronavirus twitter dataset. arxiv 2020. arXiv preprint arXiv:2003.07372.

Chen, P., Xie, H., Maslov, S., & Redner, S. (2007). Finding scientific gems with google’s pagerank algorithm. Journal of Informetrics, 1(1), 8–15.

Chen, Q., Allot, A., & Lu, Z. (2020a). Keep up with the latest coronavirus research. Nature, 579(7798), 193–193.

Chen, Q., Allot, A., & Lu, Z. (2020b, 11). LitCovid: an open database of COVID-19 literature. Nucleic Acids Research, 49(D1), D1534–D1540. doi: 10.1093/nar/gkaa952

Colavizza, G. (2020). Covid-19 research in wikipedia. Quantitative Science Studies, 1(4), 1349–1380. doi: 10.1162/qss\_a\_00080

Ghosh, R., Kuo, T.-T., Hsu, C.-N., Lin, S.-D., & Lerman, K. (2011). Time-aware ranking in dynamic citation networks. In 2011 ieee 11th international conference on data mining workshops (pp. 373–380).

Hope, T., Portenoy, J., Vasan, K., Borchardt, J., Horvitz, E., Weld, D. S., … West, J. (2020). Scisight: Combining faceted navigation and research group detection for COVID-19 exploratory scientific search. In Q. Liu & D. Schlangen (Eds.), Proceedings of the 2020 conference on empirical methods in natural language processing: System demonstrations, EMNLP 2020 - demos, online, november 16-20, 2020 (pp. 135–143). Association for Computational Linguistics. doi: 10.18653/v1/2020.emnlp-demos.18

Horbach, S. P. J. M. (2020). Pandemic publishing: Medical journals strongly speed up their publication process for covid-19. Quantitative Science Studies, 1(3), 1056–1067. doi: 10.1162/qss\_a\_00076

Kanellos, I., Vergoulis, T., Sacharidis, D., Dalamagas, T., & Vassiliou, Y. (2019). Impact-based ranking of scientific publications: a survey and experimental evaluation. IEEE Transactions on Knowledge and Data Engineering.

Kanellos, I., Vergoulis, T., Sacharidis, D., Dalamagas, T., & Vassiliou, Y. (2020). Ranking papers by their short-term scientific impact. CoRR, abs/2006.00951. Retrieved from https://arxiv.org/abs/2006.00951

Kousha, K., & Thelwall, M. (2020). Covid-19 publications: Database coverage, citations, readers, tweets, news, facebook walls, reddit posts. Quantitative Science Studies, 1(3), 1068–1091. doi: 10.1162/qss\_a\_00066

Page, L., Brin, S., Motwani, R., & Winograd, T. (1999). The pagerank citation ranking: Bringing order to the web. (Tech. Rep.). Stanford InfoLab.

Panja, S., Maan, A. K., & James, A. P. (2020). Vilokana - lightweight COVID19 document analysis. In 63rd IEEE international midwest symposium on circuits and systems, MWSCAS 2020, springfield, ma, usa, august 9-12, 2020 (pp. 500–504). IEEE. doi: 10.1109/MWSCAS48704.2020.9184598

Piwowar, H. (2013). Introduction altmetrics: What, why and where? Bulletin of the American Society for Information Science and Technology, 39(4), 8–9.

Rohatgi, S., Karishma, Z., Chhay, J., Keesara, S. R. R., Wu, J., Caragea, C., & Giles, C. L. (2020). Covidseer: Extending the CORD-19 dataset. In Doceng ‘20: ACM symposium on document engineering 2020, virtual event, ca, usa, september 29 - october 1, 2020 (pp. 21:1–21:4). ACM. doi: 10.1145/3395027.3419597

Su, D., Xu, Y., Yu, T., Siddique, F. B., Barezi, E. J., & Fung, P. (2020). Caire-covid: A question answering and query-focused multi-document summarization system for COVID-19 scholarly information management. In Proceedings of the 1st workshop on NLP for covid-19@ EMNLP 2020, online, december 2020. Association for Computational Linguistics. doi: 10.18653/v1/2020.nlpcovid19-2.14

Thelwall, M. (2020). Coronavirus research before 2020 is more relevant than ever, especially when interpreted for covid-19. Quantitative Science Studies, 1(4), 1381–1395. doi: 10.1162/qss\_a\_00083

Trewartha, A., Dagdelen, J., Huo, H., Cruse, K., Wang, Z., He, T., … Ceder, G. (2020). Covidscholar: An automated COVID-19 research aggregation and analysis platform. CoRR, abs/2012.03891. Retrieved from https://arxiv.org/abs/2012.03891

Vergoulis, T., Chatzopoulos, S., Kanellos, I., Deligiannis, P., Tryfonopoulos, C., & Dalamagas, T. (2019). Bip! finder: Facilitating scientific literature search by exploiting impact-based ranking. In Proceedings of the 28th ACM international conference on information and knowledge management, CIKM 2019, beijing, china, november 3-7, 2019 (pp. 2937–2940). ACM. doi: 10.1145/3357384.3357850

Vergoulis, T., Kanellos, I., Atzori, C., Mannocci, A., Chatzopoulos, S., La Bruzzo, S., … Manghi, P. (2021). Bip! db: A dataset of impact measures for scientific publications. arXiv preprint arXiv:2101.12001.

Vergoulis, T., Kanellos, I., Chatzopoulos, S., Pla Karidi, D., & Dalamagas, T. (2021, February). Bip4covid19: Impact metrics and indicators for coronavirus related publications. Zenodo. doi: 10.5281/zenodo.4555117

Wang, L. L., Lo, K., Chandrasekhar, Y., Reas, R., Yang, J., Eide, D., … others (2020). Cord-19: The covid-19 open research dataset. ArXiv.

Wang, Q., Li, M., Wang, X., Parulian, N., Han, G., Ma, J., … Onyshkevych, B. A. (2020). COVID-19 literature knowledge graph construction and drug repurposing report generation. CoRR, abs/2007.00576.

Wise, C., Ioannidis, V. N., Calvo, M. R., Song, X., Price, G., Kulkarni, N., … Karypis, G. (2020). COVID-19 knowledge graph: Accelerating information retrieval and discovery for scientific literature. CoRR, abs/2007.12731.

